# Behind and Beyond the Screen: Neural Differences of Live Over Video-Based Action Perception under Attentional Load

**DOI:** 10.64898/2026.02.23.707376

**Authors:** Elif Ahsen Cakmakci, Sezan Oral, Burcu A. Urgen

**Affiliations:** Department of Neuroscience, Bilkent University, Ankara, 06800, Türkiye; Department of Psychology, Bilkent University, Ankara, 06800, Türkiye; Aysel Sabuncu Brain Research Center and National Magnetic Resonance Research Center (UMRAM), Ankara, 06800, Türkiye

**Keywords:** Ecological validity, Contextual processing, Representational similarity analysis, Social presence

## Abstract

Perceiving others’ actions is essential for survival, interaction, and communication, yet most neuroscience studies rely on 2D videos or images that lack the presence and social affordances of real actions. This limits our understanding of real-world action perception and the development of neurally grounded models. Here, we directly compare behavioral and neural responses to real (live) versus video-based actions. Using a novel experimental setup (Pekçetin et al. 2023), we conducted a two-session EEG study (N = 26) in which participants viewed peripheral actions presented live or via video while performing a central task under low and high attentional load. We examined behavioral performance, mass-univariate ERPs, time-frequency responses, and time-resolved representational similarity (RSA). Behaviorally, real actions imposed a greater cognitive cost than video actions, with the largest “Realness Effect” under high load. ERPs showed reliable Live-Video differences within 150–450 ms after action onset. Time-frequency analyses over occipital and parietal regions revealed weaker alpha (8-12 Hz) and beta (15-25 Hz) suppression for video actions, indicating reduced perceptual engagement. Time-resolved RSA also robustly separated live and video conditions between 250–750 ms. Together, these results show that live actions engage perceptual systems more strongly than their video-based counterparts, underscoring the limitations of screen-mediated paradigms and motivating more ecologically grounded approaches in social and action perception research.

## 1 Introduction

### 1.1 Biological action preception

A fundamental ability shared by humans and other animals is to perceive and recognize the actions of other living beings. This capacity is vital for survival, communication, and social interaction. A key aspect of it is distinguishing biological motion, the movements produced by living organisms, from inanimate motion (Blake and Shiffrar 2007). It enables individuals to interpret others’ actions, infer meaning, and respond adaptively (Vogeley 2017). Understanding how the brain supports this capacity requires identifying the neural systems involved in processing observed actions.

Neurophysiological and neuroimaging studies, especially those using fMRI, have mapped the main brain regions involved in perceiving others’ actions. This network includes early visual areas such as V1, motion-sensitive MT+, and occipitotemporal regions like the extrastriate body area (EBA) and the posterior superior temporal sulcus (pSTS), extending into parietal and premotor cortices (Grossman and Blake 2002; Saygin 2007; Peelen and Downing 2007; Lingnau and Downing 2015; Giese and Rizzolatti 2015; Urgen et al. 2019; Urgen and Orban 2021). Within this network, early visual areas process low-level features of observed motion, MT+ is sensitive to movement, and pSTS responds strongly to biological motion, sending information to parietal and premotor areas that interpret higher-level aspects such as goals or targets (Urgen et al. 2019).

Furthermore, electroencephalography (EEG) studies examining the temporal dynamics of action recognition have shown that event-related potentials (ERPs) such as *P* 1, *N* 1, *P* 2, *N* 2, and *N* 300*/N* 400 are sensitive to biological motion (Hirai et al. 2003; Krakowski et al. 2011; Sitnikova et al. 2008; Urgen et al. 2018). Another class of EEG signals associated with action perception is cortical oscillations. During the observation of actions, significant decreases have been observed in the power of mu (8–12 Hz) and beta (15–30 Hz) frequencies recorded over electrodes positioned on the somatosensory cortex (Oberman et al. 2007; Press et al. 2011; Urgen et al. 2013). These regions have been identified in studies using both simplified point-light displays and more complex video stimuli of human actions. However, point-light displays, being visually minimal, tend to elicit weaker activation in the action perception system compared to videos (Jastorff et al. 2016). This brings up a key question: does greater stimulus realism lead to fuller engagement of the neural mechanisms that support action perception, also known as ecological validity problem. This question has recently become an emerging focus in the literature, reflecting a growing recognition that stimulus realism may critically determine how completely the neural mechanisms of action perception are engaged.

### 1.2 Previous studies on more realistic stimuli

Previous studies have attempted to increase the realism of action perception experiments by introducing stereoscopic 3D videos as an intermediate step between standard two-dimensional recordings and live presentations. This approach aimed to enhance perceptual depth and spatial immersion while retaining the controllability of video-based paradigms (Jastorff et al. 2016; Ferri et al. 2016). Findings from this line of work show that actions viewed in 3D elicit stronger activation in depth-sensitive occipitoparietal regions such as the cuneus and V7 compared to 2D viewing (Jastorff et al. 2016) and recruit additional areas beyond the canonical action perception network (Ferri et al. 2016). These include the dorsomedial parietal cortex (DIPSM), involved in depth processing and whole-body movement monitoring within peripersonal space, the ventral premotor cortex, which responds to actions directed toward the observer, and the parieto-insular cortex, a core hub of the vestibular system (zu Eulenburg et al. 2012; Frank and Greenlee 2018).

While these studies demonstrate that increasing visual realism can expand the engaged neural network, they remain limited by the inherent boundaries of screen-based presentation. Most importantly, they cannot reproduce the sense of “here and now” that arises when an observer shares the same physical space and temporal frame with the actor, an element that may be essential for fully engaging the social and sensorimotor systems involved in real-world action perception.

Only a few studies have directly compared live and video presentations of human actions (Perani et al. 2001; Järveläinen et al. 2001; Caggiano et al. 2011; Jola and Grosbras 2013; Prinsen and Alaerts 2019; Karimova et al. 2024). Together, they laid the foundation for understanding how the format of action presentation influences neural responses. Live actions have been shown to produce stronger activation in parietal areas (Perani et al. 2001) and motor cortical regions (Järveläinen et al. 2001; Caggiano et al. 2011; Jola and Grosbras 2013; Prinsen and Alaerts 2019) compared to videos, as well as greater alpha and mu suppressions (Karimova et al. 2024).

Earlier live–video comparisons established valuable groundwork but shared methodological limitations that complicate interpretation. In several designs, the actor remained visible before and after each action, creating expectation effects and baseline contamination (Karimova et al. 2024; Jola and Grosbras 2013; Caggiano et al. 2011). Other studies lacked trial-based segmentation or consistent timing cues, so differences could reflect temporal variability rather than stimulus format (Perani et al. 2001; Järveläinen et al. 2001). Spatial and depth mismatches between live and video conditions were common, with live actors typically closer than the display, reducing comparability.

Within this literature, Prinsen and Alaerts (2019) offered a notable improvement by using a voltage-sensitive shutter to restrict live-actor visibility to the trial window and by matching actor, lighting, and movements across formats. Nonetheless, their design still faced timing variability because action onset depended on the actor’s initiation after shutter opening, and spatial alignment was incomplete, as the live actor sat only 15 cm behind the shutter while the video was presented on a separate monitor at a different distance. Building on these contributions, the present study introduces a new experimental setup created in our lab that is designed to control timing, visibility, and viewing geometry in a directly comparable manner for live and video actions. Details are provided in the Methodology section.

### 1.3 Attentional Mechanisms and Biological Action Perception

Most previous research on action perception has focused on selective attention, where participants actively attend to the observed actions. However, a full understanding of natural, ecologically valid action perception requires considering the actions that are incidental to the observer’s primary goal i.e., actions that happen to occur in the environment without being the explicit focus of attention, rather than merely being task-irrelevant. A few studies have examined this process. Thornton and Vuong (2004) showed that peripheral biological motion distractors can interfere with task performance even when ignored, indicating automatic processing. This research is foundational because it demonstrates that the perceptual system is constantly engaged by incidental information in the periphery, even when selective attention is devoted elsewhere. The ability of these incidental actions to compete for processing resources is a crucial aspect of real-world perception. Supporting this, Saygin and Sereno (2008) found with fMRI that unattended actions still evoke activity in occipitotemporal regions, though parietal and premotor response decrease when attention is diverted. Later, Nizamoglu and Urgen (2024) extended this work by manipulating attentional load during a central task and showed that the processing of unattended actions depends on overall attentional demands. The work of Nizamoglu and Urgen (2024) further highlights the importance of studying incidental actions, showing that their automatic processing is not absolute but is flexibly constrained by available cognitive resources. This suggests that real-world processing of co-present agents reflects an integration of bottom-up action-related signals with top-down attentional control. The findings of Nizamoglu and Urgen (2024) are closely related to perceptual and attentional load theory (Lavie 1995, 2005), which proposes that perceptual capacity is limited and that the processing of task-irrelevant stimuli depends on the demands of the primary task. When attentional load is low, residual capacity allows peripheral or unattended stimuli to be processed, whereas under high load, most resources are allocated to the central task, reducing or even eliminating such processing. In line with this framework, Nizamoglu and Urgen (2024) showed that neural resources to unattended biological motion decrease as attentional demands increase, indicating that the extent of automatic action processing is constrained by available perceptual capacity. Building on this framework, the present study examines how attentional load interacts with the realism of observed actions, comparing neural responses to live and video stimuli. By presenting live and video actions as incidental events in the periphery, this study provides a high-fidelity model for investigating the continuous monitoring of the environment, a mechanism vital for everyday social interactions that cannot be captured by paradigms focused purely on selective attention.

## 2 Methods

### 2.1 Task and Stimuli

All stimuli were programmed in Python (*v*3.8) using PsychoPy (*v*2023.2.3). The task is inspired by the paradigm of Schwartz et al. (2005), where the participants focus on a stream of centrally presented “T” letters and respond to specific targets while ignoring peripheral visual events appearing on either side of the screen. Each trial comprised eight “T” stimuli that varied in color (red, yellow, green, blue) and orientation (upright or inverted). Two attentional-load conditions were used. In the low-load condition, participants pressed the spacebar for any red “T.” In the high-load condition, they responded only to upright yellow and inverted green “T”s, which required attention to both color and orientation (Figure 1 (a)). Each “T” subtended roughly 1.35*° ×* 1.9*°* of visual angle and was displayed for 500 ms, separated by a gray screen of 250–350 ms. Each sequence contained two target events, one in the first 3 seconds and one in the last 3 seconds, ensuring that attention to the central task was sustained throughout the trial (Figure 1 (b)).

**Fig. 1.**
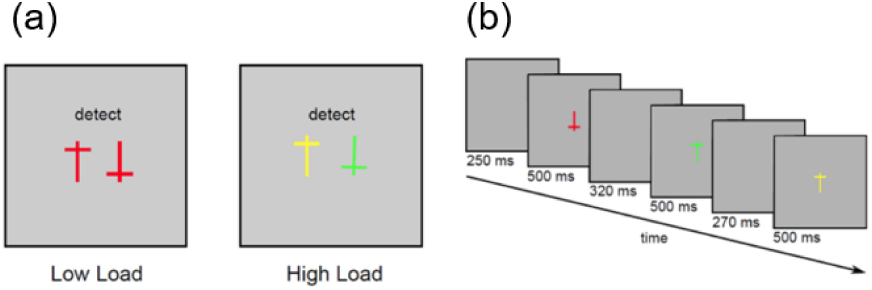
(a) Target “T” stimuli in the experimental task based on attentional load manipulation. (b) Exemplary temporal sequence of the central stimuli.

Peripheral events consisted of short human actions presented either live or as pre-recorded videos. Each stimulus lasted six seconds. The actor remained still for the first three seconds, followed by the execution of one of six actions during the final three seconds: throwing an object, extending a hand for a handshake, waving, reaching, walking, or scratching the head. The initial static period was included for two reasons. First, in the real condition, the visibility of the actor was controlled using LED lighting behind the screen, and introducing motion only after a stationary interval prevented potential interference from light-related transients that were not present in the video condition. Second, this structure allowed the onset of the action itself, rather than stimulus appearance, to be used as the temporal reference for EEG analyses, with the third second marking action onset. Actions were presented to the left or right of the central task at approximately 7.3*°* of visual angle. This structure preserved the natural timing of movements and avoided abrupt onsets that could distort EEG responses. A visual control condition was also included where no peripheral stimuli were shown. For real-time trials, the actors stayed out of visual angle behind the drapes. Participants were aware that an actor was present in the room, but the screen remained static throughout the trial. This condition served as a baseline for assessing whether the real and video presentations differed in any way beyond their visual properties like social presence.

The present study builds on these earlier efforts by introducing a controlled presentation system developed in our laboratory, adapted from the design proposed by Pekçetin et al. (2023), which allows real and video actions to be displayed within the same physical frame and spatial configuration where both presentation modes used the same Planar^®^ LookThru^TM^ OLED screen. In the live condition, the screen became transparent when backlit, revealing the actor behind it; in the video condition, the same screen displayed the recording of that actor performing the identical action under matched spatial layout.

Lighting and screen transitions were computer-controlled for precise synchronization with trial timing. Between trials, gray screens lasting 5–5.2 seconds allowed actors to reposition, and the same intervals were used in the video sessions for temporal consistency.

The experiment room was divided into two sections by the semi-transparent OLED screen and black drapes surrounding it. The participant area section consisted of a chair, table, keyboard, chinrest, and EEG setup. The stage area section behind the screen included a waiting zone with two stools that was not visible to the participants. Actors sat there between trials when they were not performing any actions. The backstage also contained a prompter computer that displayed action instructions to the actors, a baby monitor with night-vision capability used by the experimenter to ensure that everything proceeded correctly, and marked areas indicating where each action should be performed. The stage background was covered with a gray drape that served as a neutral background (Figure 2).

**Fig. 2.**
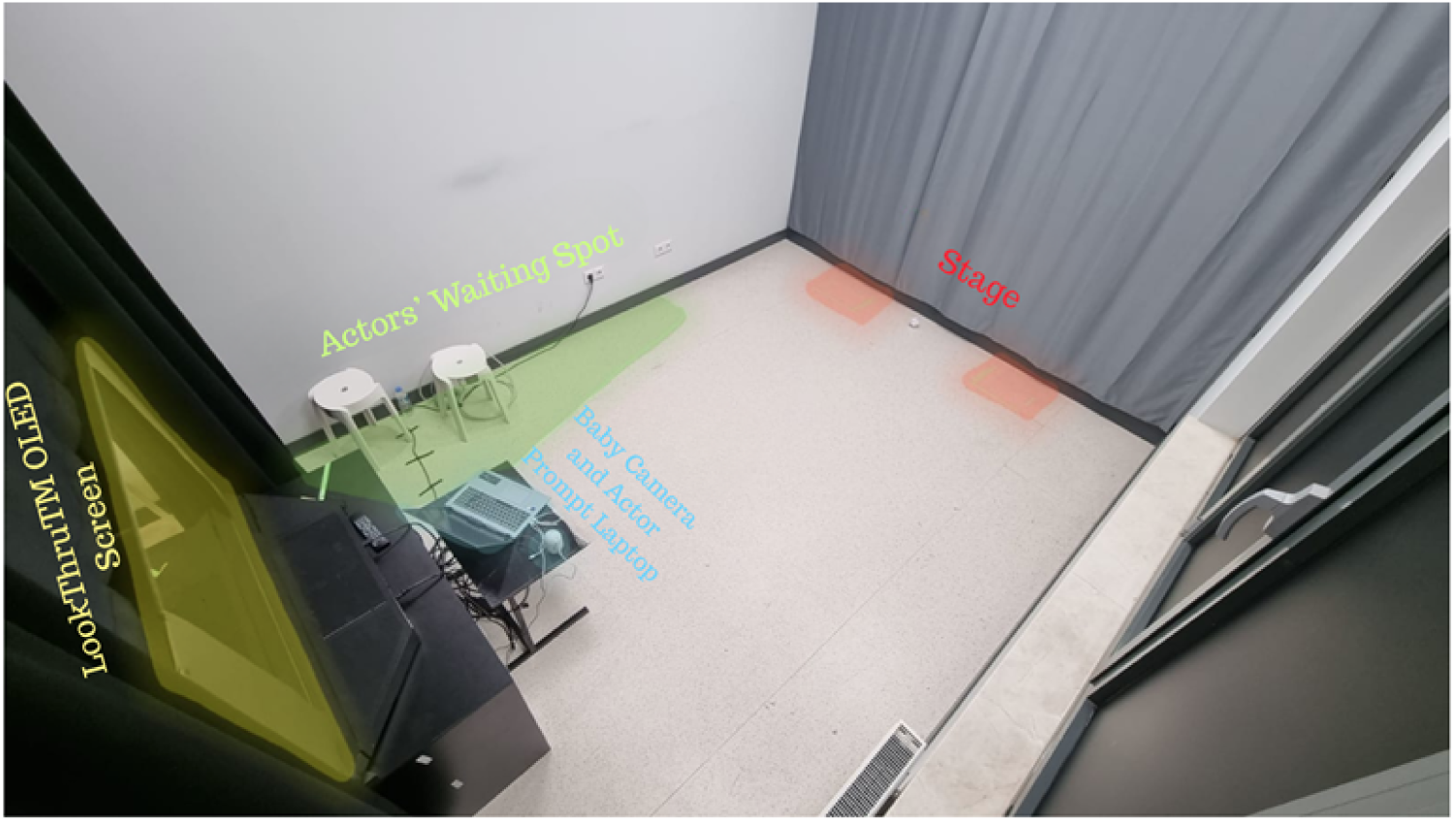
Backstage area and descriptions.

### 2.2 Experimental Design and Procedure

Each participant completed two sessions: one in which peripheral stimuli were presented as videos and another in which the same actions were performed live by actors. The order of sessions was counterbalanced across participants. Each session consisted of 16 blocks (eight high-load and eight low-load). The order of high-load and low-load blocks was randomized in pairs for each participant. The six peripheral actions were shown across all combinations of actor identity (female, male), hand used (left, right), visual field (left, right), and attentional load (high, low). Each combination was repeated twice, resulting in 192 action trials per session. A corresponding set of no-action trials was included, yielding 384 trials in total. Trial sequences were pseudo-randomized within each session. Each trial lasted about 6 seconds, followed by a gray screen of 5–5.2 seconds. At the beginning of each block, participants viewed a brief instruction screen describing the central task. Including short breaks, 384 trials, each session lasted approximately 1.5 hours. Participants responded to the central task by pressing the spacebar on a standard keyboard. They were instructed to focus on the central “T” task and to ignore the peripheral stimuli entirely.

EEG signals were recorded using a 64-channel BrainVision ActiCAP system with sintered Ag/AgCl active wet electrodes. Data were sampled at 5 kHz with 0.1*µV* resolution, and electrode impedances were maintained below 18*k*Ω.

### 2.3 Participants

Thirty-two participants were initially recruited. Data from three pilot subjects and three participants with excessive EEG noise were excluded, leaving 26 participants (9 female, 17 male; age range: 19–33 years, mean = 22.3). Half of the participants completed the live session first, and the rest began with the video session. All participants had normal or corrected-to-normal vision and were right-handed. Before the main experiment, they completed a short training session using video stimuli and were required to achieve at least an 80% F1 score in both attentional load conditions.

### 2.4 Data Preprocessing and Analysis

For behavioral analysis, task performance was quantified using the F1 score. The F1 score was computed for each condition and entered into a 2 (Attention Load: low, high) × 2 (Stimulus Type: live, video) × 2 (Action Type: action, no action) repeated-measures ANOVA. Post hoc comparisons were performed when significant interaction effects were found (Bonferroni correction, *q < .*05).

EEG data were processed in MATLAB using the EEGLAB toolbox (version 2024.2) (Delorme and Makeig 2004). Signals were band-pass filtered between 1 and 40 Hz and re-referenced to the common average. Channels with poor signal quality were visually identified and interpolated (a maximum of 10 per participant). Artifacts from eye movements and blinks were removed using Independent Component Analysis (ICA), with up to four components rejected per participant.

For ERP analysis, data were epoched from *−*500 to +1000 ms relative to the onset of the peripheral action (*t* = 0, corresponding to the third second of each trial). For time–frequency (ERSP) analyses, epochs extended from *−*1000 to +2500 ms. Noisy trials were rejected through visual inspection, and at least 30 clean trials per condition were required.

Event-related potentials (ERPs) were averaged within each of the eight experimental conditions (2 × 2 × 2 design). Statistical analysis was performed using the ffdrGND function from the Factorial Mass Univariate ERP Toolbox (version 0.5.1) (Groppe et al. 2011), which applies FDR correction across time and electrode dimensions (Benjamini–Hochberg correction, *q < .*05).

To assess the representational structure of neural responses, Representational Similarity Analysis (RSA) was performed using the rsatoolbox in Python (version 0.3.0) (Nili et al. 2014). Representational dissimilarity matrices (RDMs) were computed using correlation distance across the 0–1000 ms post-onset window. Time-resolved RSA examined the evolution of similarity patterns in 50 ms intervals, and multidimensional scaling (MDS) was applied to visualize condition relationships over time.

Time–frequency representations were computed using complex Morlet wavelets implemented in FieldTrip (version 20240111) (Oostenveld et al. 2011). Oscillatory power was estimated from 2–40 Hz (1-Hz resolution) over a *−*500 to +2500 ms window surrounding the onset of the peripheral action. A 6-cycle wavelet was used across all frequencies, with temporal estimates obtained every 20 ms. Power values were baseline-normalized using the *−*500 to 0 ms interval preceding the action onset and converted to decibel (dB) units, yielding event-related spectral perturbation (ERSP) maps for each trial.

Group-level analyses were conducted using Monte-Carlo cluster-based permutation tests implemented in FieldTrip. For each main effect and interaction in the 2 *×* 2 *×* 2 design, subject-level ERSP difference maps were compared using dependent-samples t-statistics, with cluster-based correction applied across the time–frequency plane (8–30 Hz, 0–2000 ms). All tests used 1000 permutations and a two-tailed cluster-forming threshold of *p < .*05. This procedure was used for all contrasts in the 2 *×* 2 *×* 2 design, including both main effects and interactions.

## 3 Results

### 3.1 Behavioral

A three-way ANOVA on F1 scores with factors Session (live, video), Load (high, low), and Distractor (action, no action) revealed significant main effects of Session and Load. Participants performed more accurately in the video session (*M* = 0.753, *SD* = 0.3) than in the live session (*M* = 0.693, *SD* = 0.337), F(1, 21443) = 214.25, *p < .*001, *ηp*2 = 0.01 and under low load (*M* = 0.815, *SD* = 0.291) compared to high load (*M* = 0.631, *SD* = 0.322), *F* (1, 21443) = 1963.04, *p < .*001, *ηp*2 = 0.084 (Figure 3).

**Fig. 3.**
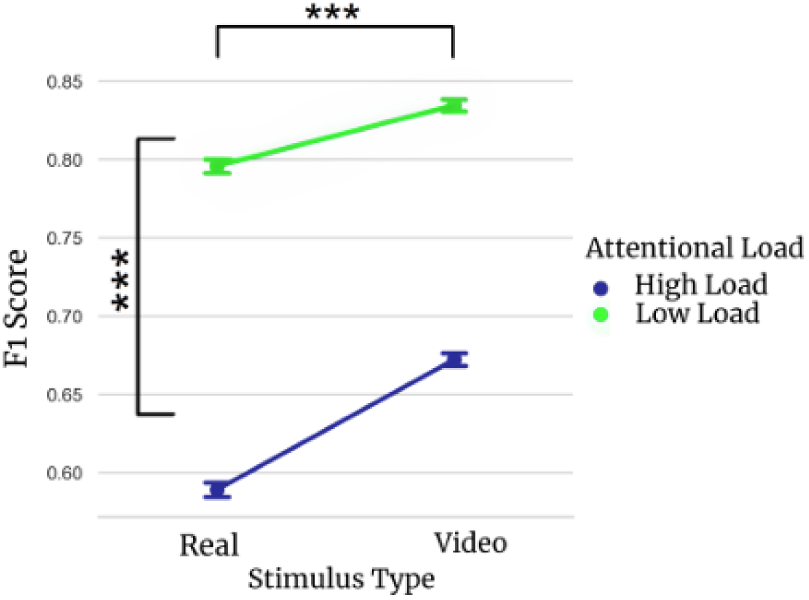
Mean F1 scores by attentional load and stimulus type. Error bars represent standard error.

A significant Session *×* Load interaction was found, *F* (1, 21443) = 28.16, *p < .*001, *ηp*2 = 0.001 indicating that the effect of attentional load was stronger in the live session. The main effect of Distractor (action vs. no action) and all its interactions were not significant (all *p > .*05).

### 3.2 ERP Results

ERP analyses revealed significant main effects of Session (live vs. video) and Attentional Load (high vs. low).

The main effect of Load showed widespread temporal and spatial distribution, with enhanced negative and positive peaks for high-load trials compared to low-load trials, consistent with greater attentional demand. (Figures 4-6)

**Fig. 4.**
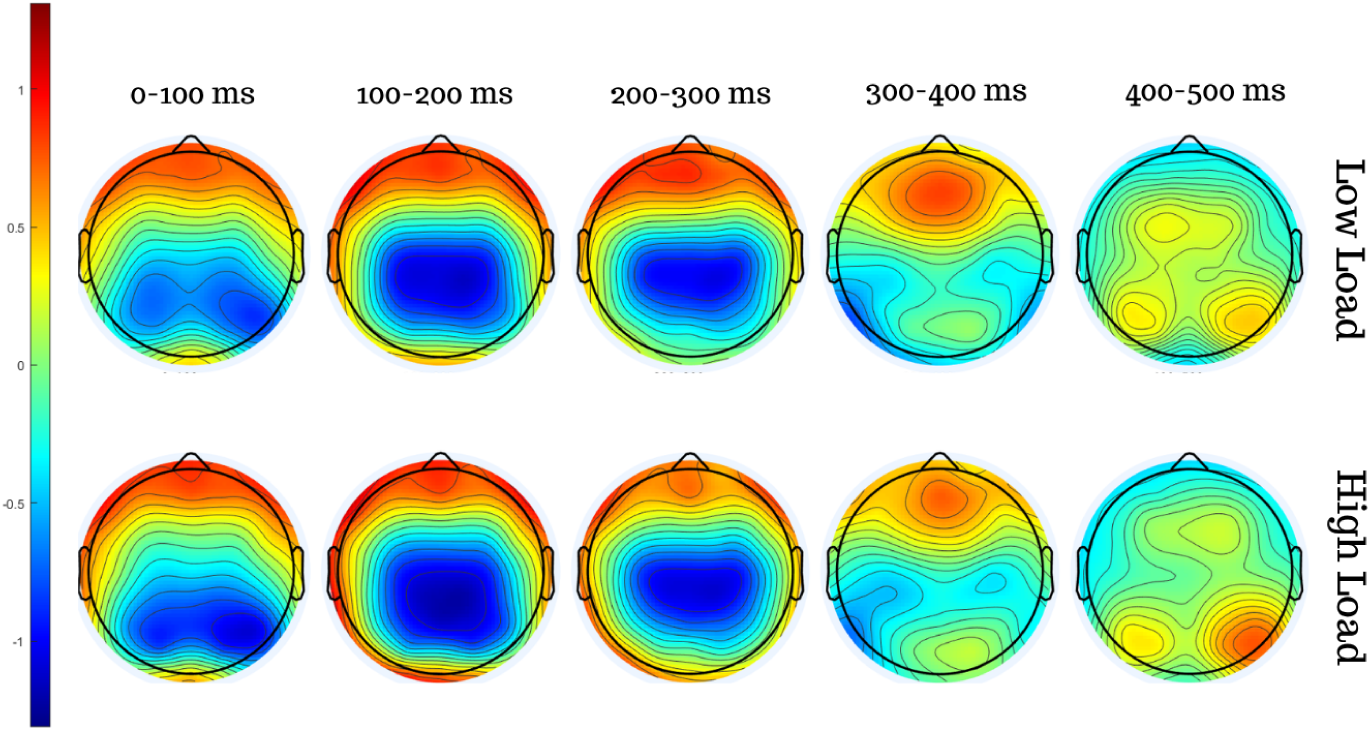
Topographic maps showing differences between Low Load and High Load conditions for the 0-500 ms time window with 100 ms steps.

**Fig. 5.**
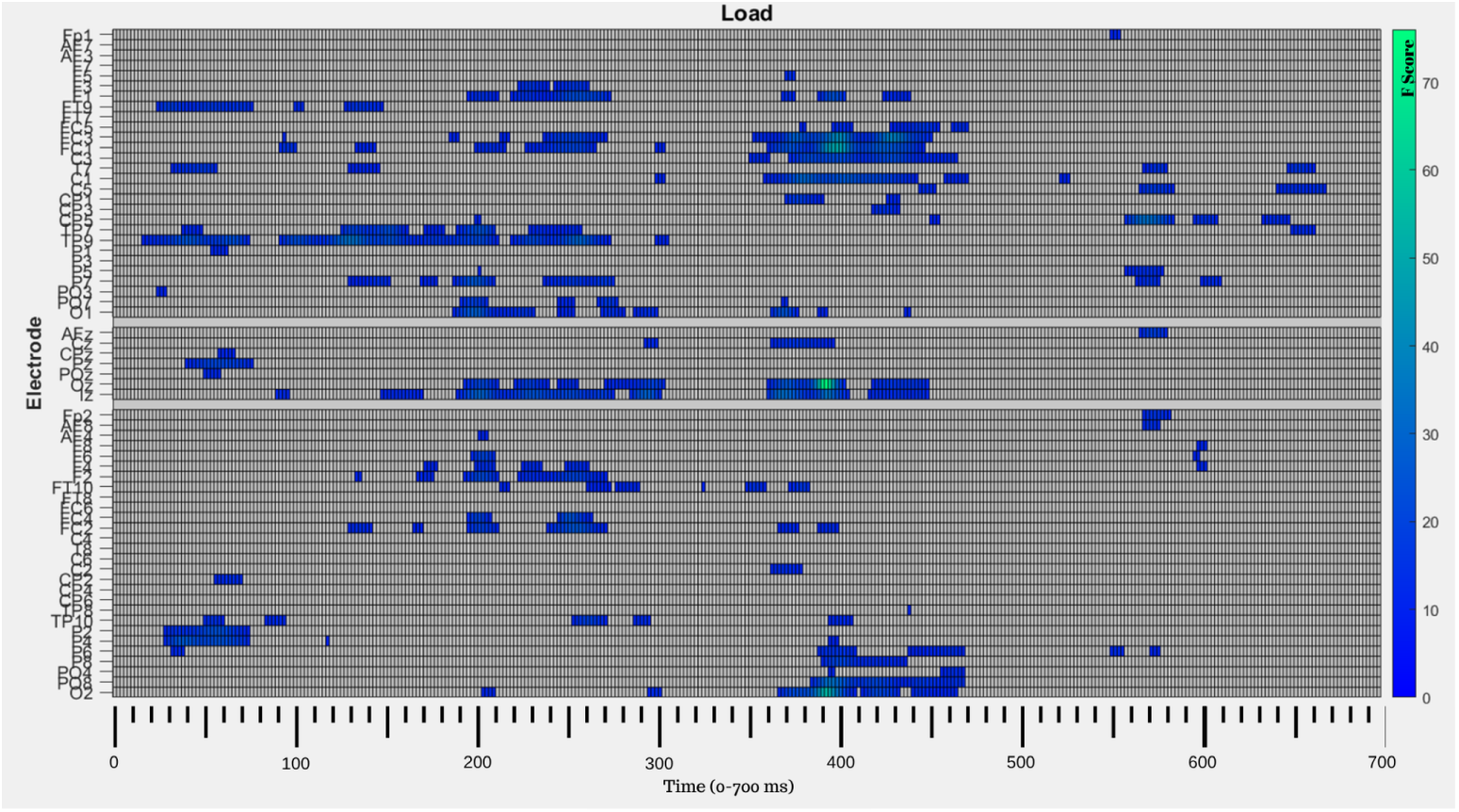
ANOVA results for the main effect of Load, the channel, and time windows where the effect was significant (*p <* 0.05) are marked in color, where the shade of color varies according to the F-score values.

**Fig. 6.**
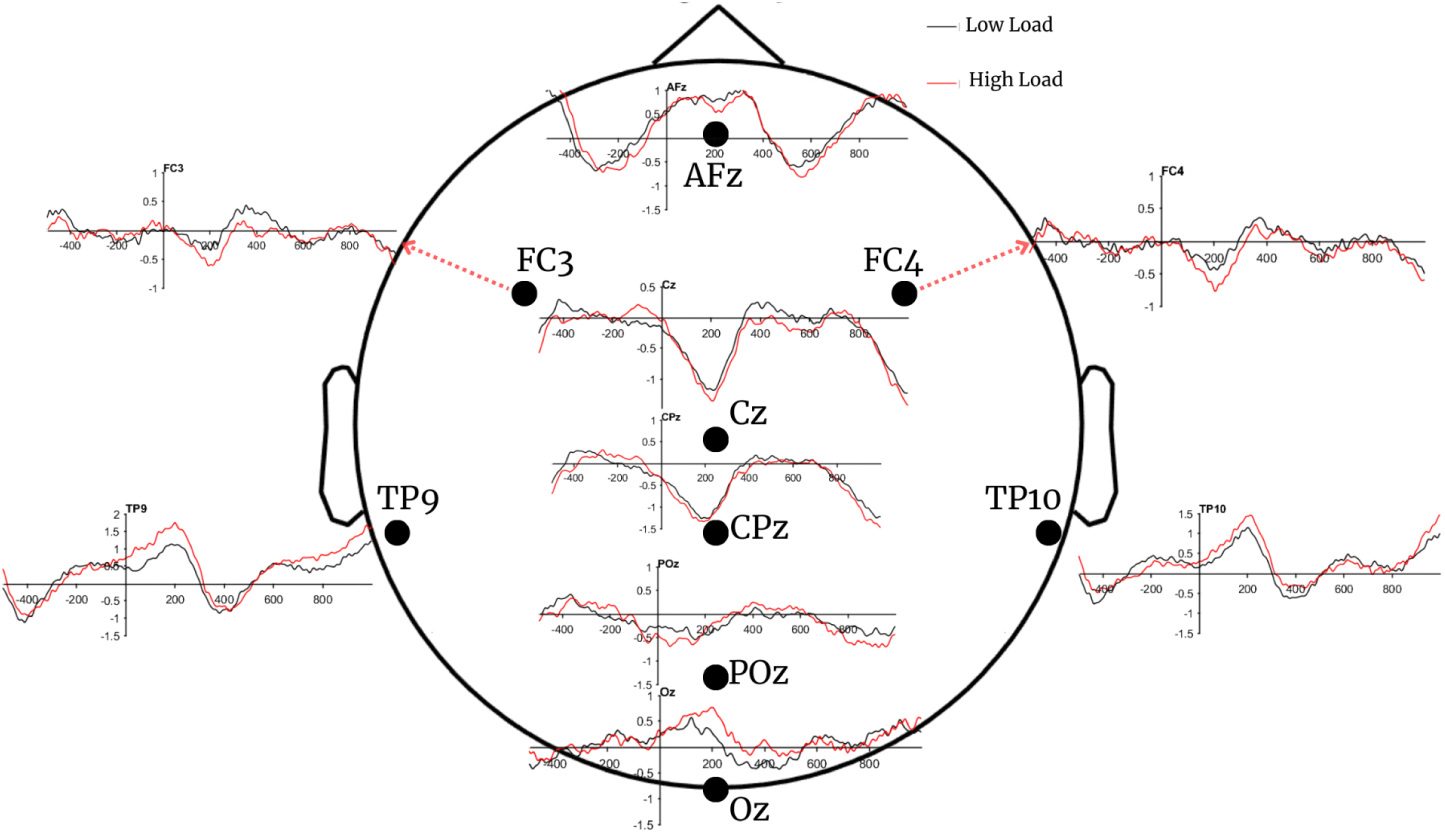
Example waveforms from electrodes showing significant differences between Low Load and High Load conditions.

The main effect of the Session showed widespread spatial distribution too and was most pronounced within the 250–450 ms time window. (Figures 7-9)

**Fig. 7.**
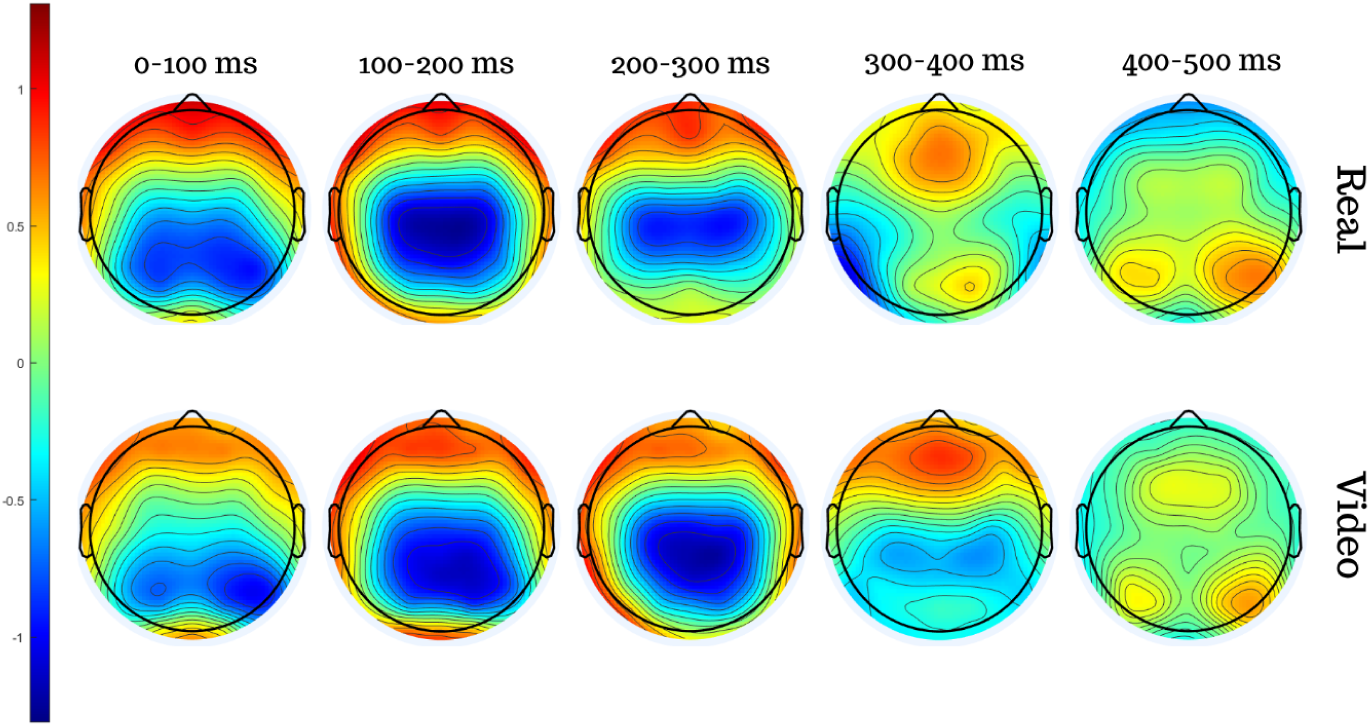
Topographic maps showing differences between Video and Real Stimuli conditions for the 0-500 ms time window with 100 ms steps.

**Fig. 8.**
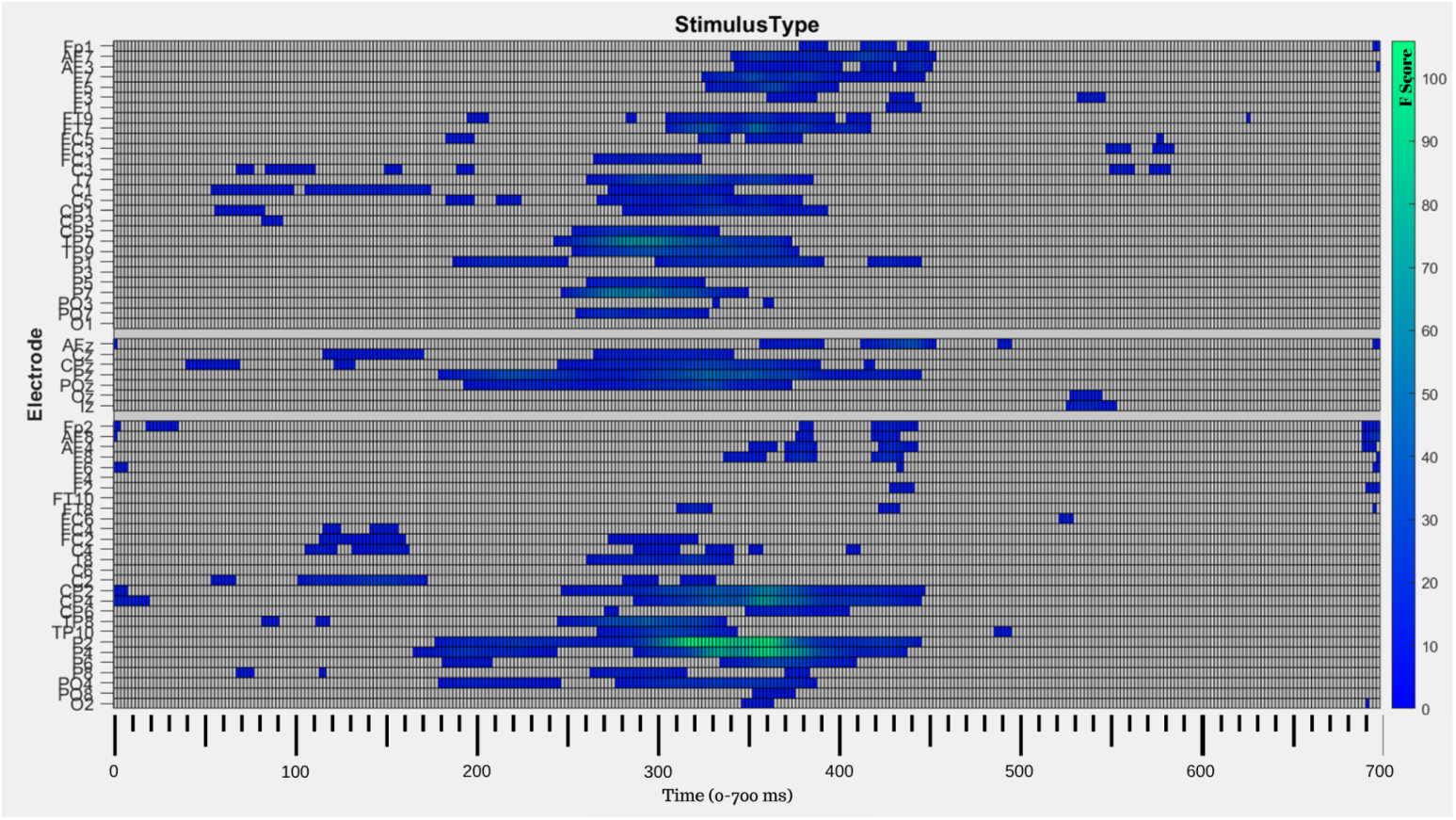
ANOVA results for the main effect of Stimulus Type, the channel, and time windows where the effect was significant (*p <* 0.05) are marked in color, where the shade of color varies according to the F-score values.

**Fig. 9.**
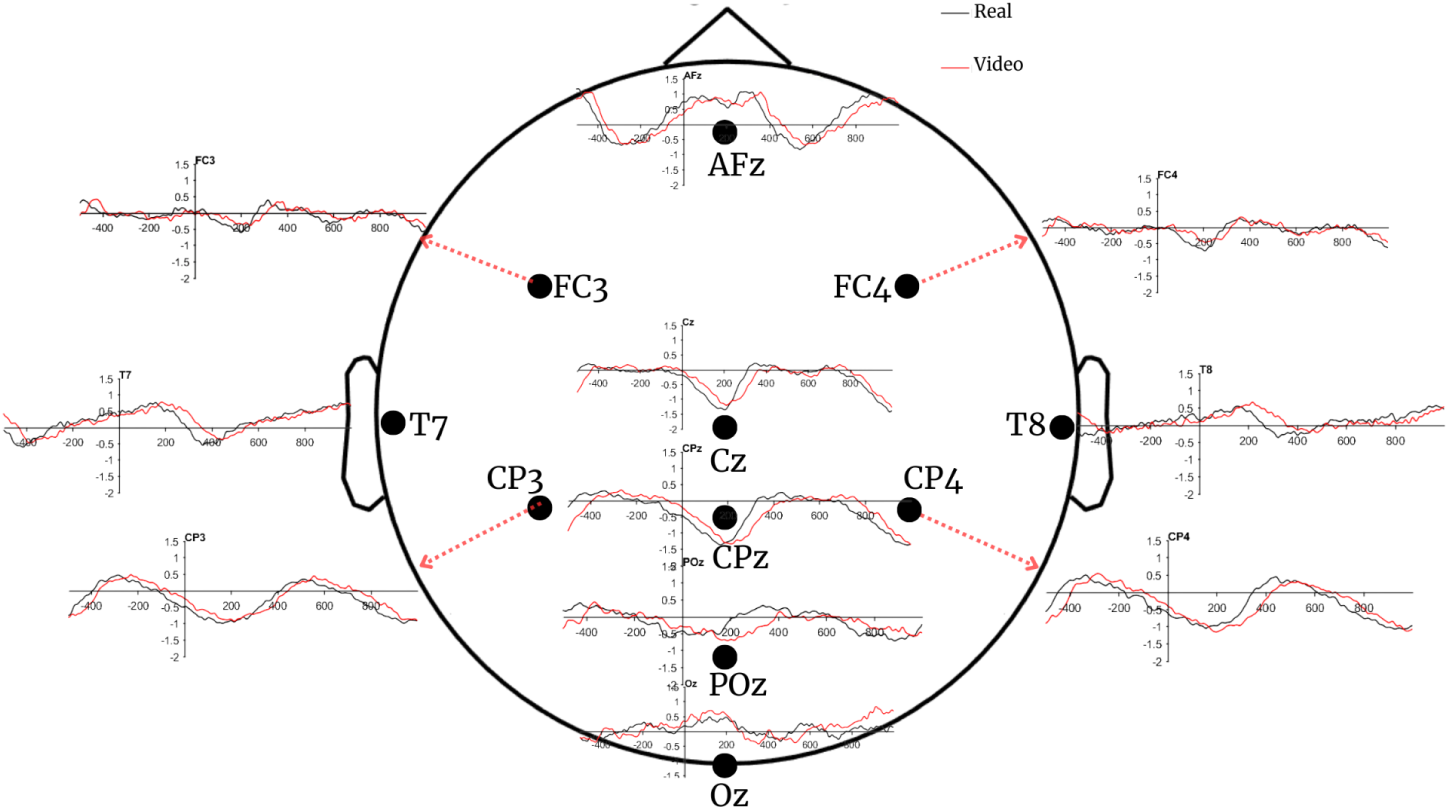
Example waveforms from electrodes showing significant differences between Real and Video Stimuli conditions.

No other main effects or interactions reached significance after correction for multiple comparisons.

### 3.3 RSA

The representational dissimilarity matrix (RDM) computed from the group-averaged EEG data in the 0–1000 ms window (Figure 10) revealed that the clearest neural separation occurred between live and video conditions. Within this structure, live trials showed greater dissimilarity among themselves than video trials, suggesting that neural responses to live actions were more variable across attentional load and action type.

**Fig. 10.**
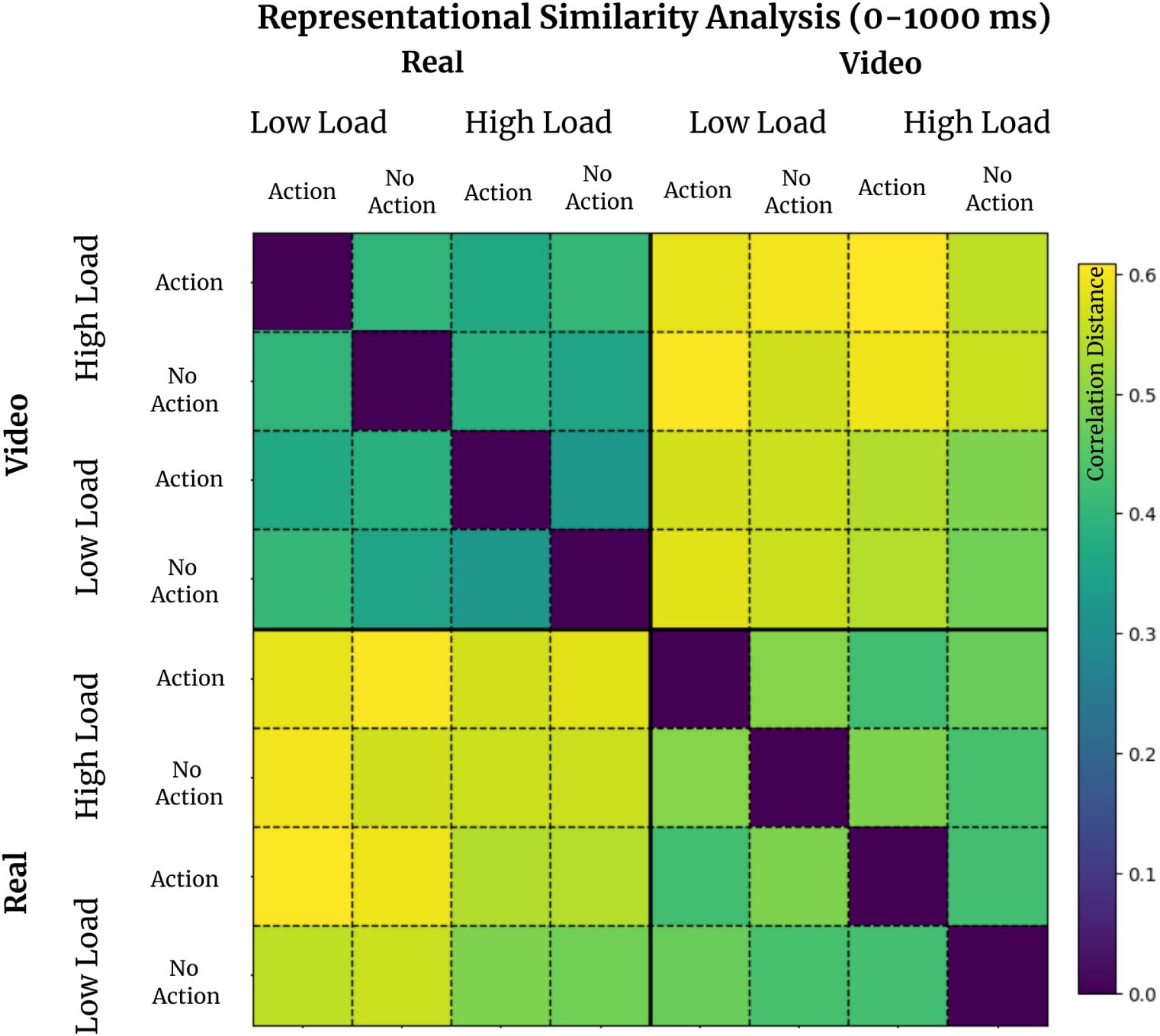
The Representational Dissimilarity Matrix (RDM) illustrates the correlation–based dissimilarity of multichannel ERP patterns between each pair of experimental conditions across the 0–1000 ms time window. Because correlation distance (1–*ρ*) was used as the distance metric, yellow tones indicate greater dissimilarity, while purple tones indicate higher similarity.

Temporal RSA results (Figure 11) showed that condition-related dissimilarities began to emerge around 250 ms and reached their peak between roughly 550 and 700 ms. During this period, the most pronounced divergence appeared between live and video conditions, consistent with the pattern in the static RDM. Also, the earliest occurrence and the latest decay of differentiation were between video and live conditions. After approximately 800 ms, representational distances gradually decreased, indicating a convergence of neural patterns across conditions.

**Fig. 11.**
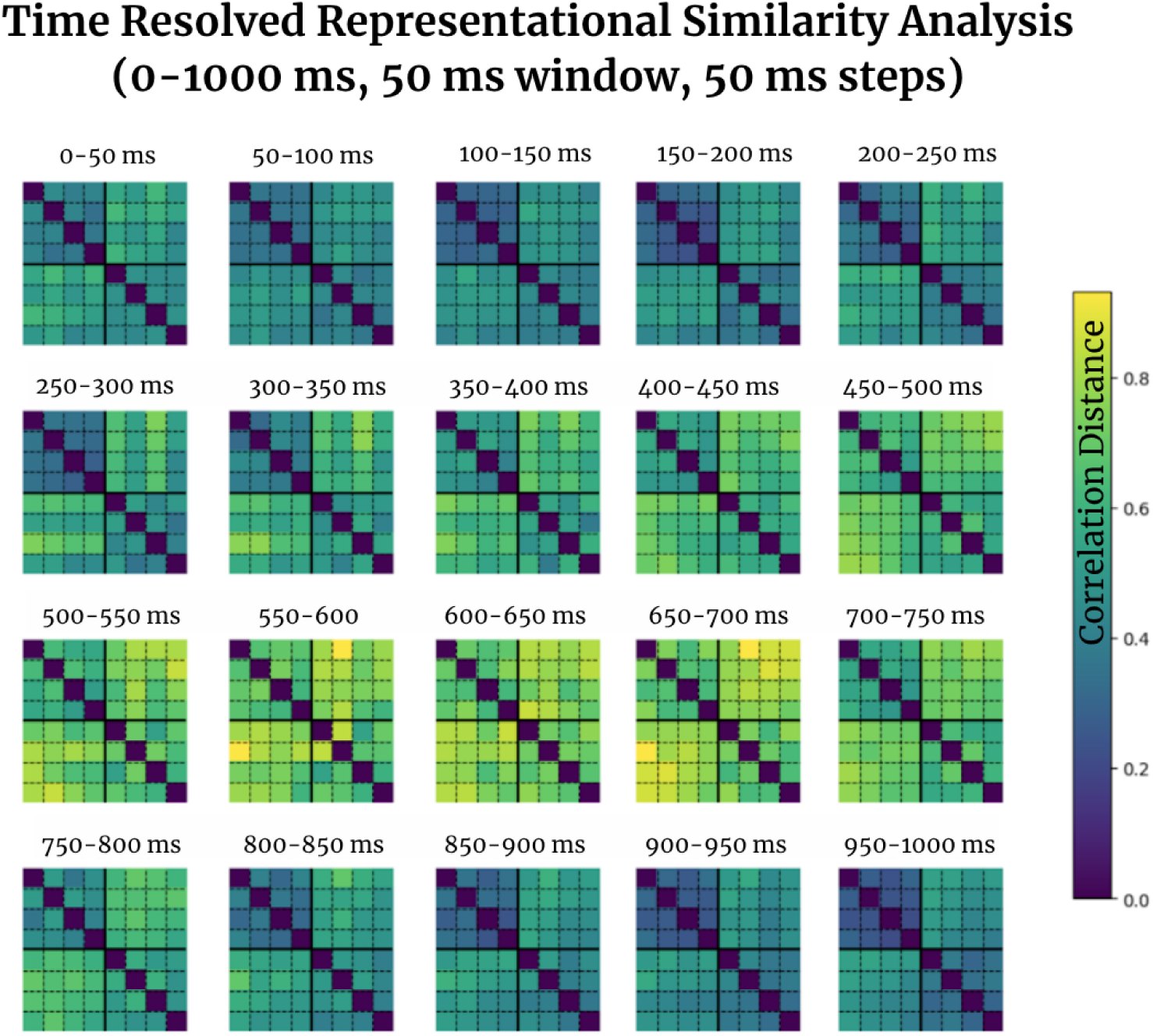
Temporally resolved RDM, computed in 50 ms steps across the 0–1000 ms time window.

The multidimensional scaling (MDS) projection derived from the static RDM supported this pattern (Figure 12). Points corresponding to live and video conditions formed two clearly separated clusters, suggesting that stimulus format rather than attentional load or action type primarily structured the neural representational geometry across the 0–1000 ms window.

**Fig. 12.**
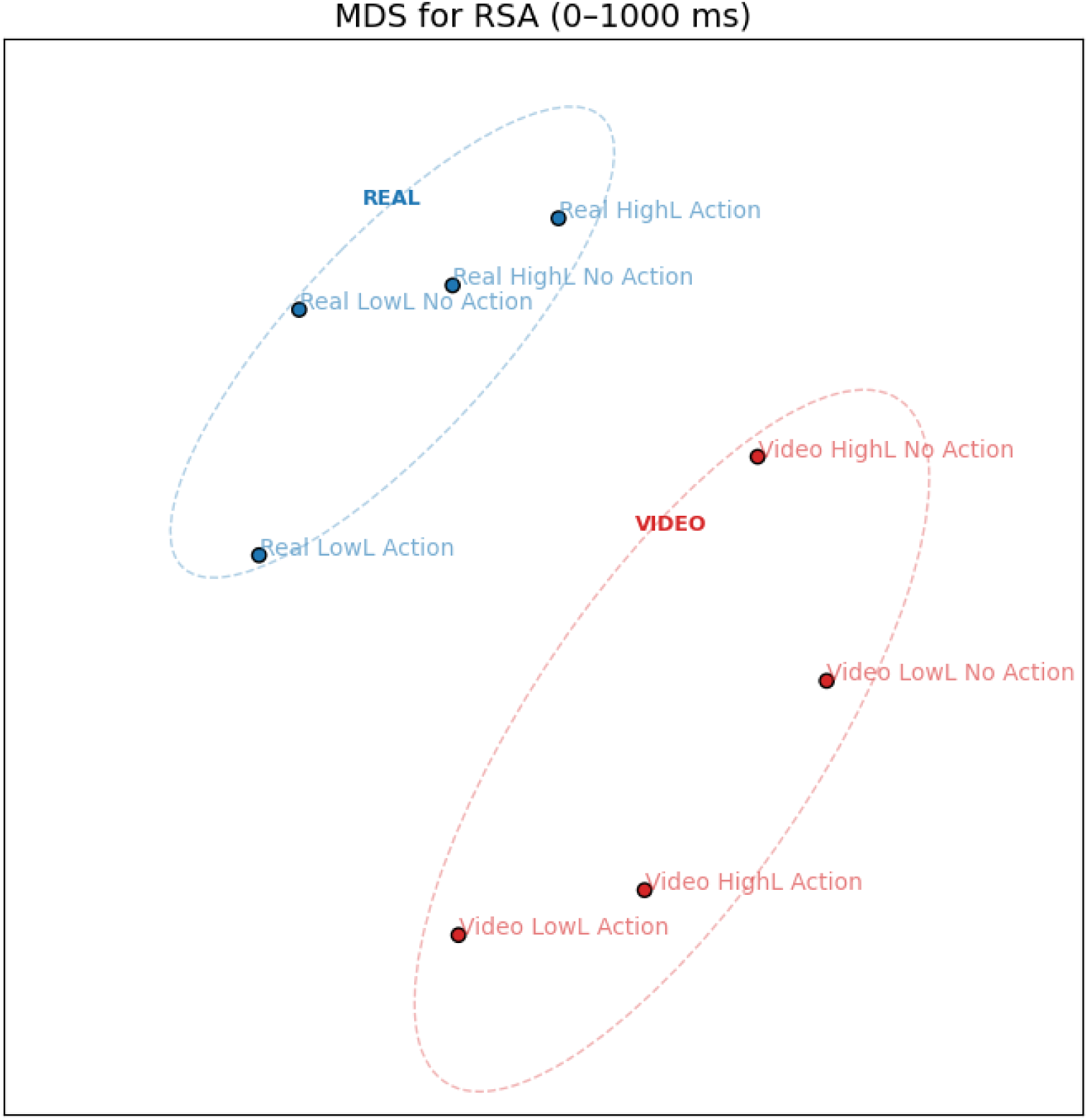
The MDS visualization summarizes the neural representations of experimental conditions in a two-dimensional space based on the RDM calculated over the 0–1000 ms time window. The MDS visualization summarizes the neural representations of experimental conditions in a two-dimensional space based on the RDM calculated over the 0–1000 ms time window.

### 3.4 Time-Frequency Analysis

Statistical analysis on ERSP data showed a main effect for all three factors: Load (High and Low Load), Action (Action and No Action), and Stimulus (Real and Video Stimulus). The load main effect was prominent over the parietal and occipitoparietal electrodes, with more suppression for Low Load in the Alpha and Beta bands. (Figure 13)

**Fig. 13.**
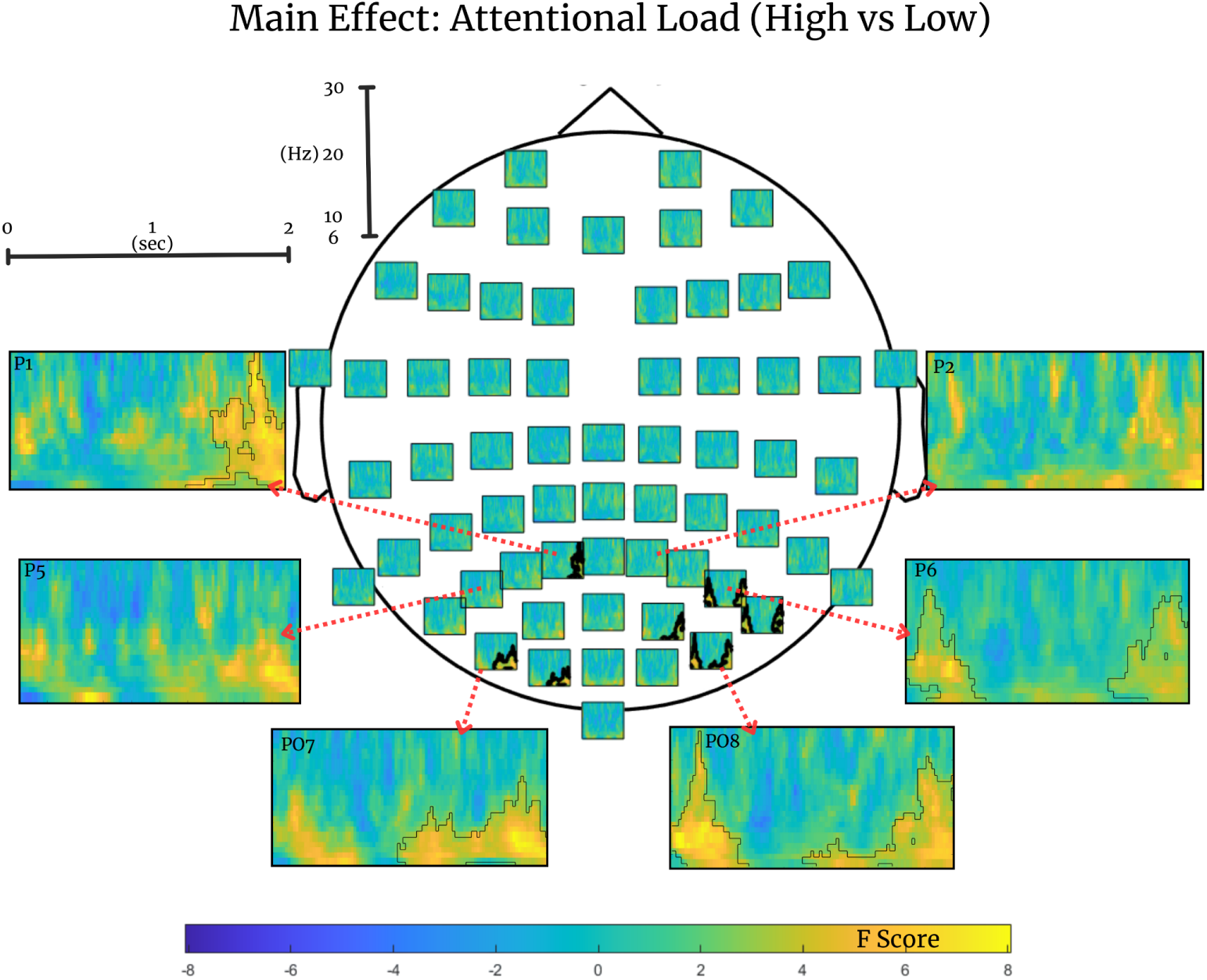
ERSP difference maps from electrodes showing significant differences (marked with black borders) between High and Low Load conditions, where the x-axis is time (0 *−* 2 sec) and the y-axis is the frequency (6 *−* 30 Hz) interval.

The main effect of Action was prominent over the parietal, occipitoparietal, central, and left frontal electrodes, with more suppression for Action in the Alpha, Mu, and Beta bands. (Figure 14)

**Fig. 14.**
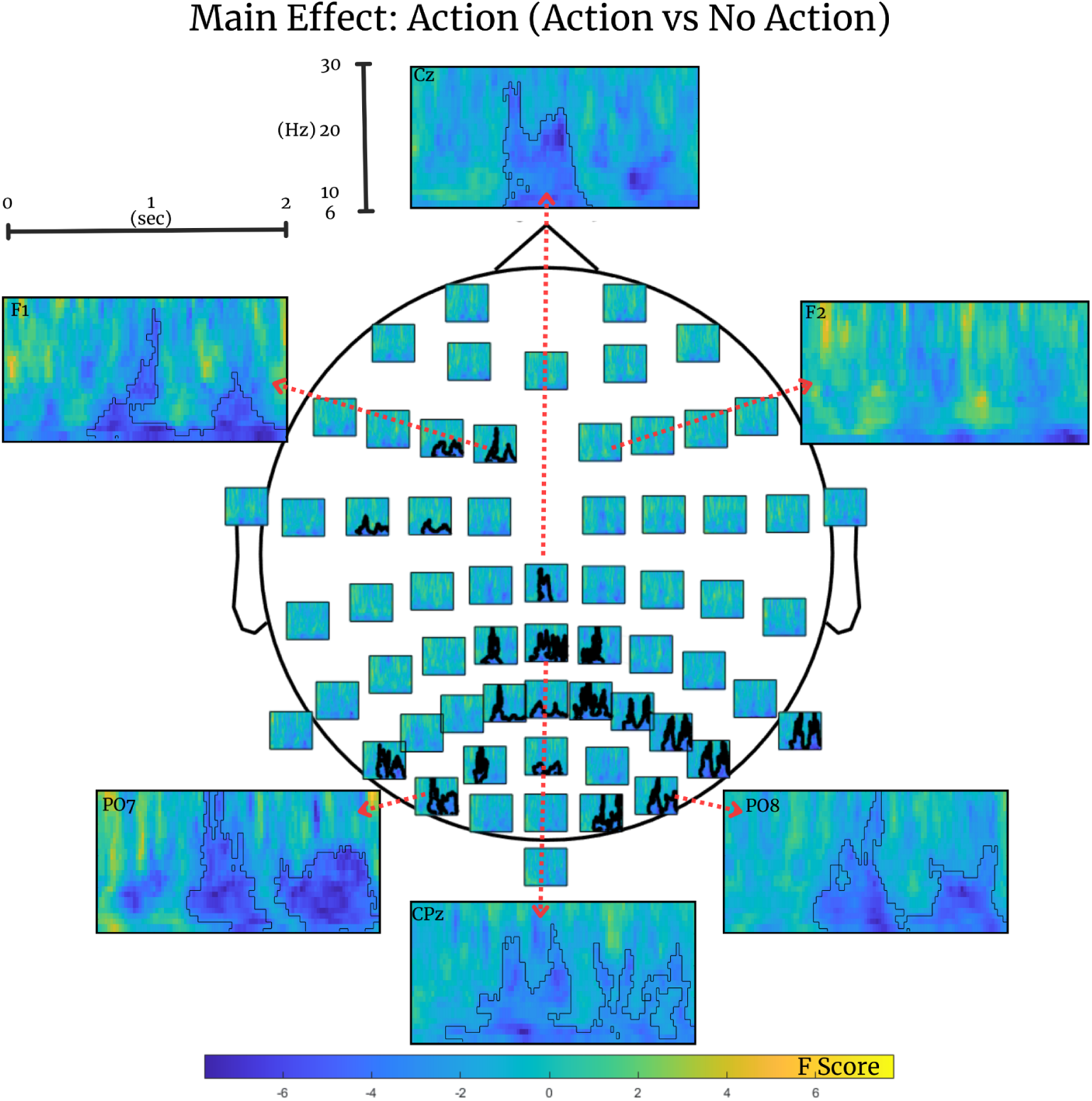
ERSP difference maps from electrodes showing significant differences (marked with black borders) between Action and No Action conditions, where the x-axis is time (0 *−* 2 sec) and the y-is the frequency (6 *−* 30 Hz) interval.

The main effect of Stimulus was prominent over the parietal and occipitoparietal electrodes, with more suppression for Real Stimulus in the Alpha and Beta bands. (Figure 15)

**Fig. 15.**
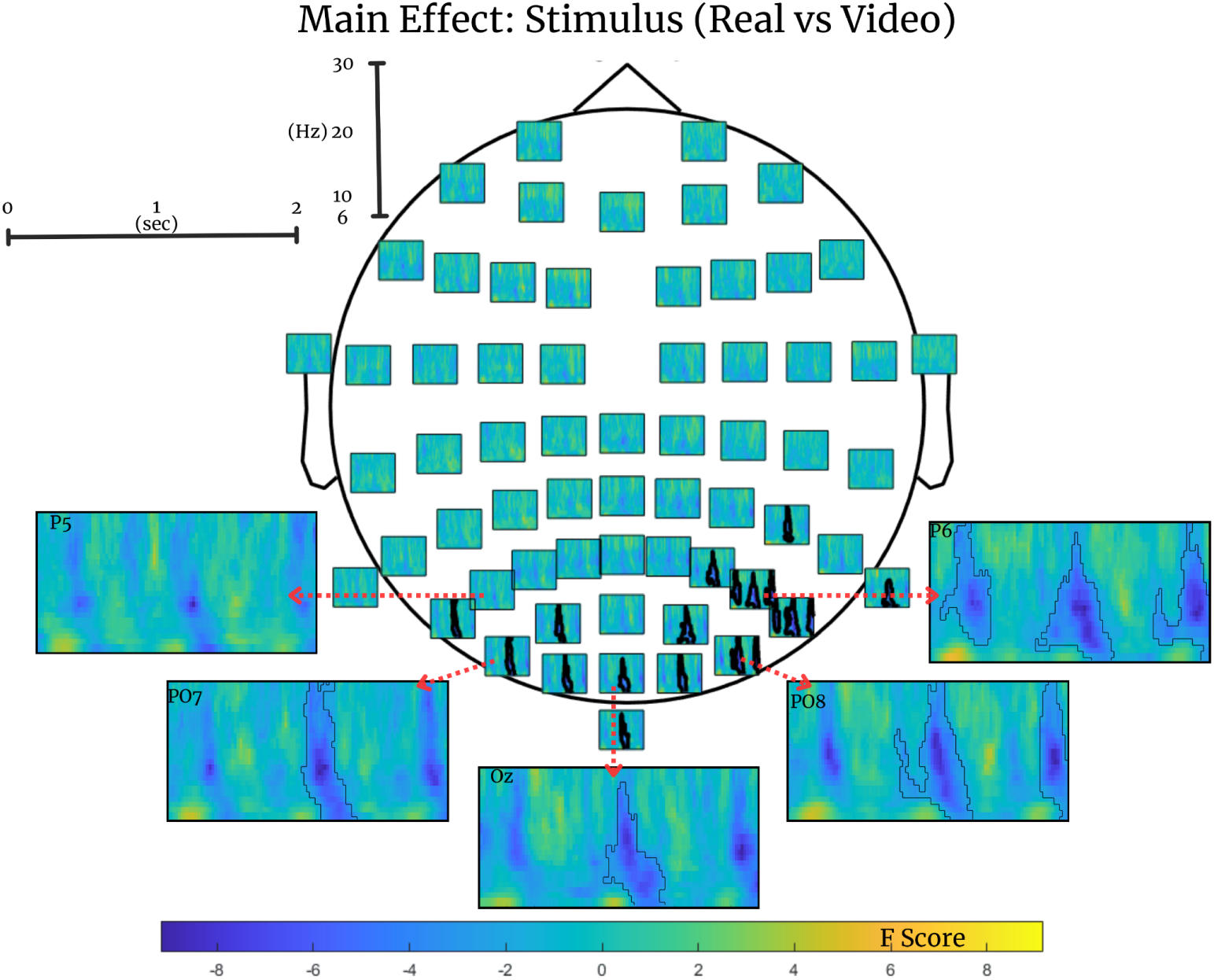
ERSP difference maps from electrodes showing significant differences (marked with black borders) between Real and Video Stimuli conditions, where the x-axis is time (0 *−* 2 sec), and the y-axis is the frequency (6 *−* 30 Hz) interval.

A spatially isolated Stimulus *×* Action interaction emerged, reaching significance at two non-adjacent electrodes (F1 and POz) (see Supplementary Figures 1-2 for ERSP interaction map and post hoc ERSP difference maps). Given its restricted topography and atypical distribution, this effect should be interpreted with caution. No other statistically significant interaction was found.

## 4 Discussion

### 4.1 Real versus video as the dominant factor across measures

Across behavioral and neural analyses, stimulus format (real vs. video) emerged as the most dominant factor. Behaviorally, performance was consistently lower for real stimuli than for video stimuli, indicating that the real condition imposed a stronger overall interference than the video condition. Neural measures mirrored this dominance. ERP analyses revealed a broad and sustained main effect of stimulus format, peaking around 250-450 ms and extending across much of the scalp. The widespread nature of this effect suggests that real and video stimuli are differentiated at multiple processing levels, reflecting a global evaluation of the scene rather than a localized sensory response.

Time–frequency analyses further supported this pattern. Real stimuli elicited stronger alpha and beta suppression over parietal, occipitoparietal, and occipital scalp regions, indicating stronger engagement of visual and visuospatial processing systems compared to video stimuli. Representational similarity analysis most clearly emphasized the dominance of stimulus format. Neural patterns were primarily organized according to whether stimuli were real or video, rather than by attentional load or action type. Across the 0-1000 ms window, real and video conditions showed clear separation, with representational differences emerging around 250 ms and peaking between approximately 550 and 700 ms. Multidimensional scaling further revealed two well-separated clusters corresponding to real and video stimuli, with minimal separation based on load or action. Notably, real trials exhibited greater dissimilarity among themselves than video trials. This suggests that neural responses to real stimuli may vary more across load and action contexts, potentially reflecting the richer or more dynamic processing demands associated with perceiving a real human in the environment.

### 4.2 Beyond visual features: physical presence and agency

Importantly, the real vs. video difference was not limited to action trials. Performance differences were also observed in no-action trials, where no visible movement occurred. In these trials, participants could not see the actor behind the scenes, but they were aware that the actor was physically present in the room. This pattern demonstrates that the behavioral difference between real and video conditions cannot be explained solely by visible peripheral motion. This aligns with Distraction-Conflict Theory (Baron 1986), which suggests the presence of others creates a conflict between task-focus and environmental monitoring. Furthermore, as noted by Markus (1978), the mere presence of another person can impair performance on cognitively demanding tasks, even without explicit evaluation.

Neural measures converged with this behavioral finding. ERP analyses revealed that the real-video distinction persisted in no-action trials, despite the absence of overt movement. The presence of a reliable stimulus-format effect under these conditions indicates that ERP differences were not driven by visual motion cues, but rather by the broader context associated with the real versus video condition.

Oscillatory results further support this interpretation. Real–video differences in alpha and beta bands were observed across analyses, indicating stronger engagement of visual and visuospatial processing systems for real stimuli. The persistence of these differences across conditions suggests that oscillatory activity was sensitive to stimulus format itself, rather than being contingent on the presence of visible action. Together, the continuation of real–video differentiation in no-action trials across behavioral performance, ERPs, and oscillatory measures indicates that features beyond visual motion contribute to the observed effects. Awareness of physical presence appears sufficient to modulate both task performance and neural processing, implying that real stimuli introduce a distinct contextual factor that is not captured by video presentations.

### 4.3 Interdependence of Physical Presence with Cognitive Load and Peripheral Action

The interaction between stimulus format and attentional load observed for behavioral results represents an additional aspect of the findings. The amplification of load-related performance costs in the real condition indicates that stimulus format shapes how task demands manifest behaviorally. When cognitive resources are already taxed by a task, the “distraction cost” of a real human presence is more pronounced than that of a video. Moreover, a significant yet limited interaction between Stimulus (Real-Video) and Action (Action-No Action) was observed. Although not dominant across measures, this interaction points to an additional dimension of processing in which action-related information is modulated by stimulus format. This pattern suggests that action perception may not operate independently of the broader stimulus context, but instead reflects an integration of action-related signals with properties associated with real versus video presentation. While the present results do not allow a detailed characterization of this interaction, its emergence is nevertheless interesting. The fact that an interaction is detectable, even under conditions where actions are peripheral and secondary, indicates that stimulus format can influence how action-related information is processed at a fundamental level. This raises the possibility that under paradigms where actions are more central to the scene and more strongly attended, such effects may become more pronounced and structured, revealing additional aspects of how real and video stimuli shape action perception.

### 4.4 Context-Dependent Anticipation and its Reflection in Event-Related Potentials

The emergence of a significant ERP main effect for stimulus format, in the absence of a corresponding action effect, warrants further investigation. While the gradual nature of biological motion may have been insufficient to elicit a transient evoked response, the consistent, time-locked differentiation between real and video formats suggests that this neural activity is specifically tied to the anticipated action phase of the trial. This differentiation likely arises because participants internalized the trial structure, where a static period is followed by a possible movement. Through repetition, this pattern becomes predictable, and the transition point at which movement could occur triggers a shift in context processing-even if no movement actually follows. Because a real human body carries greater social or behavioral relevance than a video image, this potential for action generates a stronger, time-locked neural response in the Real condition. Furthermore, the presence of this Real-Video effect in No-Action trials confirms that the differentiation is not driven by visual motion cues, but rather by a state of readiness modulated by physical presence.

### 4.5 Attentional load effects across measures

Attentional load exerted a robust influence on behavioral performance. Participants performed worse under high load, confirming that the central task effectively taxed attentional resources. This decrease in performance indicates that fewer resources were available for handling competing information when task demands increased.

At the neural level, attentional load produced widespread ERP amplitude differences across the time course and scalp. Because ERPs were time-locked to the onset of the peripheral action rather than the central task stimuli, these effects are expected. The load manipulation was present throughout the trial and could therefore influence perceptual processing, decision-related activity, and motor preparation simultaneously, rather than being confined to a specific processing stage.

Time–frequency analyses further reflected the impact of attentional load. Alpha and beta power differed reliably between low- and high-load conditions over parietal and occipitoparietal regions. Stronger alpha and beta suppression in the low-load condition is consistent with greater engagement of visual and visuospatial processing systems when cognitive demands from the central task are reduced. When attentional resources are not fully consumed, the perceptual system appears more responsive to peripheral input. Together, these findings indicate that attentional load acts as a global factor that shapes behavioral performance and neural activity across multiple levels of analysis.

### 4.6 Action versus no-action effects across measures: convergences and dissociations

In contrast to attentional load, action-related effects showed a more differentiated pattern across measures. Behaviorally, no reliable main effect of action was observed, indicating that the mere presence of peripheral movement did not substantially alter task performance. ERP analyses likewise revealed no robust differentiation between action and no-action trials. This limited ERP sensitivity to action content can be explained by the temporal structure of the experiment. ERPs were time-locked to the onset of the peripheral action, which occurred several seconds after the initial static scene, making early visual components that typically distinguish action-related stimuli less likely to be captured. Moreover, the gradual onset of biological motion, rather than an abrupt visual change, may have further reduced sensitivity relative to classical ERP paradigms. In contrast, time-frequency analyses revealed a clear main effect of action, with enhanced mu suppression reflecting increased sensorimotor engagement. Unlike ERPs, oscillatory measures are less dependent on precise temporal onsets, making them more sensitive to sustained or gradually emerging action-related processing.

Time-frequency analysis showed differences between action and no-action trials across the alpha, mu, and beta bands over parietal, parieto-occipital, centroparietal, and left frontal electrodes. Stronger alpha and beta suppression in action trials likely reflects increased visual and visuospatial engagement when peripheral actions are present. Enhanced mu suppression suggests greater sensorimotor engagement, consistent with the involvement of motor systems during the observation or anticipation of human actions. Additional suppression over left frontal regions may reflect increased involvement of action monitoring, prediction, or executive control processes. The left-lateralized pattern may partly relate to the predominantly right-handed sample, as the actions were performed with both hands and presented on both sides of the periphery. Right-handed individuals often show stronger left-hemisphere recruitment during action-related processing (Cabinio et al., 2010); thus, this asymmetry likely reflects participant neurobiology rather than stimulus properties. Taken together, these results indicate that action-related information is more readily captured by oscillatory measures than by ERPs or behavioral measures, highlighting the complementary sensitivity of different analytical approaches.

### 4.7 Limitations and Future Work

Despite these contributions, the present study represents a step toward, rather than a full realization of, naturalistic experimentation. Real-world perception involves many factors that cannot be controlled simultaneously, and here we deliberately adopted an intermediate approach that allowed a well-matched comparison between real and video stimuli while minimizing major confounds related to size, timing, spatial layout, and stimulus identity. Nevertheless, some differences between the two conditions remain unavoidable. In the real condition, illumination was provided by ambient ceiling lights, whereas in the video condition, the primary light source was the display itself, and luminance levels were not explicitly equated across formats. In addition, although stimulus size was carefully matched, the real actor was positioned behind the screen while the video was presented on the screen surface, potentially introducing subtle differences in perceived distance. More broadly, the controlled nature of the setup may have constrained the sense of realness: participants and actors were not allowed to interact, and the transparent screen, while enabling precise control, likely limited available action affordances. Future studies employing more interactive and flexible setups may therefore reveal additional aspects of real action perception that could not be captured here. Moreover, the present design raises several open questions. The analyses were time-locked to action onset; aligning neural responses to stimulus appearance instead may reveal different temporal dynamics or interaction patterns. Similarly, presenting actions at fixation rather than in the periphery may alter the balance between real and video processing, potentially amplifying or reducing the observed differences. Addressing these questions will be critical for further disentangling the roles of presence, attention, and action in real-world perception.

## 5 Conclusion

This study examined neural and behavioral differences between real and video peripheral stimuli, with low and high attentional load imposed by a central task. By directly comparing live and video stimuli within a tightly controlled yet naturalistic setup, the present work addressed a central limitation of the existing literature: the assumption that visual approximation implies perceptual approximation. Across behavioral and neural measures, this assumption was not supported.

Behavioral results showed that real peripheral stimuli disrupted central task performance more strongly than video stimuli, an effect that persisted even in the absence of visible peripheral distractors. The performance decrement observed in real no peripheral stimulus trials (No Action) demonstrates that the influence of stimulus format cannot be reduced to visual features alone. Instead, the results point to a broader contextual factor, namely the awareness of a physically co-present agent, which alters attentional allocation even when no action is being performed. This finding suggests that perceptual systems remain sensitive to the possibility of action, not only to its visual occurrence. Also, the interaction between attentional load and stimulus type showcases that the attentional load change affects participants more during real stimuli presentation. Taken together, the results demonstrate that perception of real stimuli is meaningfully different from its video counterpart, indicating that there are mechanisms that cannot be fully understood by solely using video stimuli.

Neurally, this distinction was reflected across multiple analysis domains. ERP results revealed reliable and widespread differences between real and video conditions emerging at mid-latencies and extending across the scalp. These effects were present regardless of whether a peripheral action occurred, supporting the conclusion that stimulus format itself modulates neural processing.

Time–frequency analyses further showed that real stimuli elicited stronger suppression in alpha and beta bands over parietal and occipitoparietal regions compared to video stimuli, indicating enhanced engagement of visuospatial processing systems when peripheral events were physically present. In contrast, mu-band suppression was specifically associated with the presence of an action, distinguishing action from no-action trials rather than real from video contexts. Representational similarity analysis converged with these findings by showing that neural activity patterns were primarily organized by stimulus format. Real and video conditions formed clearly separable representational states over time, while distinctions related to attentional load or action presence played a secondary role.

Collectively, the findings underscore the importance of ecological validity in studies of action perception and attention. Real stimuli are not simply more vivid versions of videos; they reorganize attentional priorities and neural processing by embedding perception within a physically and socially grounded context. To understand how the brain processes actions as they occur in everyday life, experimental paradigms must account for presence, not only presentation. Moving toward such paradigms is essential for bridging the gap between laboratory-based models of perception and real-world cognition.

## Data Availability

Raw EEG data and data analysis codes used in this study can be accessed through OSF: https://osf.io/qtdjk/ upon request.

## Appendix A

**Fig. A1.**
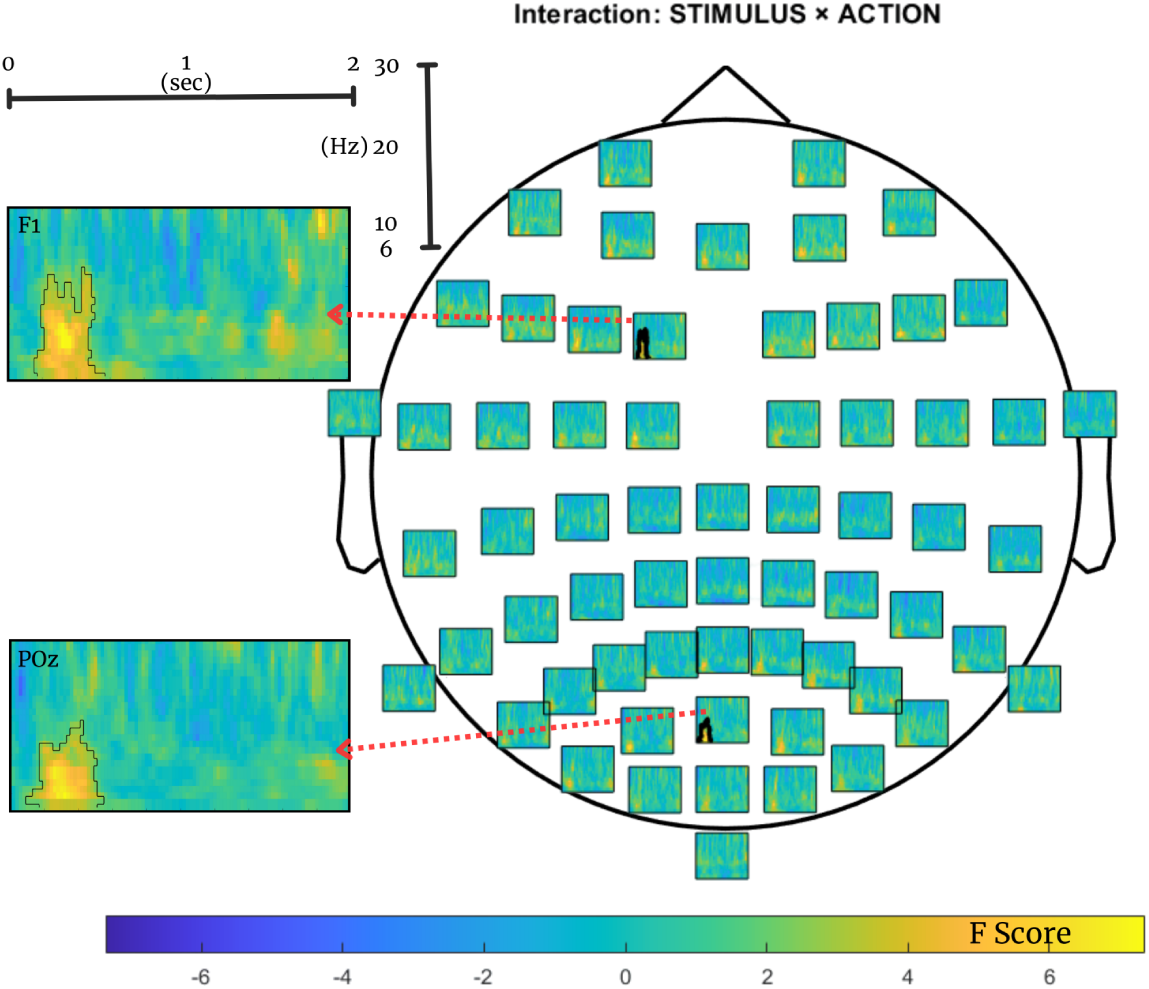
ERSP difference maps from electrodes showing significant interaction (marked with black borders) between Action and Stimulus Type features, where the x-axis is time (0-2 sec) and the y-axis is the frequency (6-30 Hz) interval.

**Fig. A2.**
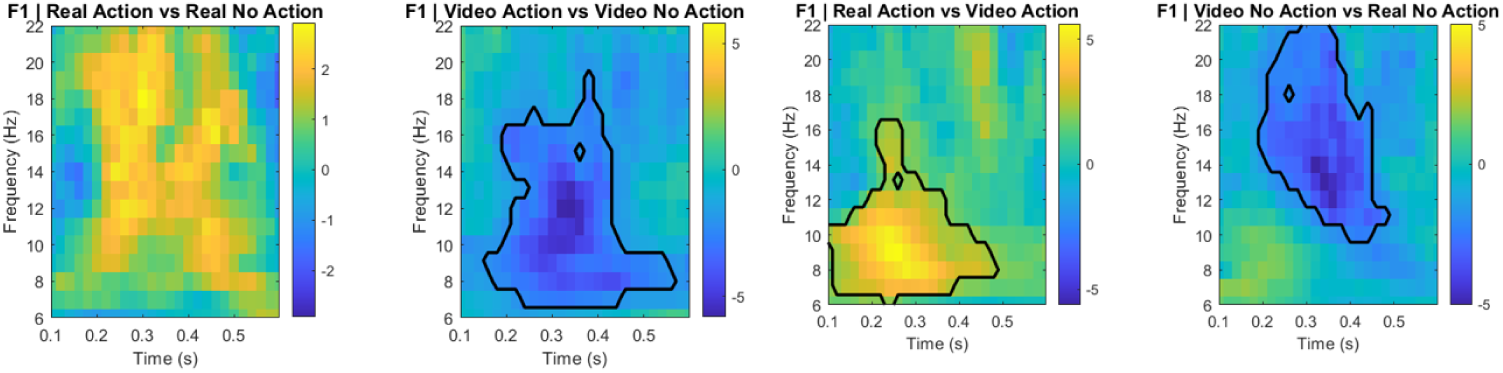
Post hoc plots for Action and Stimulus Type interaction for F1 electrode where the x-axis is time (0.1-0.6 sec), and the y-axis is the frequency (6-22 Hz) interval and significant differences are marked with a black border.

**Fig. A3.**
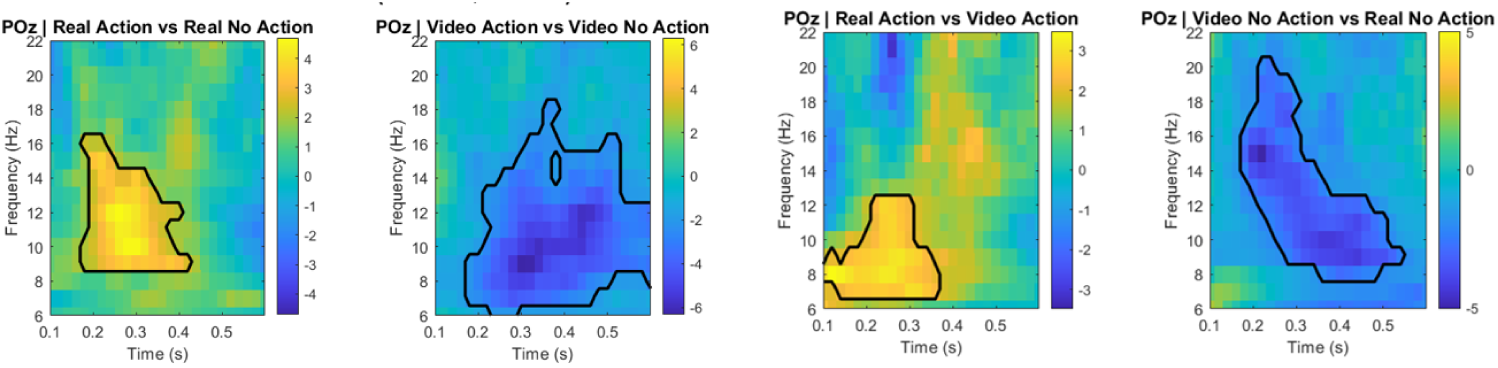
Post hoc plots for Action and Stimulus Type interaction for POz electrode, where the x-axis is time (0.1-0.6 sec), and the y-axis is the frequency (6-22 Hz) interval, and significant differences are marked with a black border.

## Acknowledgements

As authors, we wish to express deepest gratitude to Zeynep Nese Bardakcı and Emirhan Hızoglu for their roles as actors, Papatya Yesilbas, Can Atasayar and Tuvana Karaduman for their support during data collection.

## Funding

This study was funded by a TUBITAK (The Scientific and Technological Research Council of Türkiye) grant, under the 1001 program (Project No: 122K915), awarded to Dr. Burcu Ayșen Ürgen.

## Contributions

EAC: Data recordings and preparation, formal analysis, investigation, methodology, visualization, writing, review and editing. SO: Conceptualization, data recordings and preparation, formal analysis, investigation, methodology, visualization, writing, review and editing. BAU: Conceptualization, funding acquisition, investigation, methodology, project administration, supervision, review and editing.

## Ethics declarations

### Conflict of interest

The authors declare no competing interests.

### Ethical approval

The study was approved by the Human Research Ethics Committee of Bilkent University.

### Consent to participate

Informed consent was obtained from all individual participants included in the study.

## Notes

https://osf.io/qtdjk/

### Competing Interest Statement

The authors have declared no competing interest.

